# Identification of differentially methylated single-nucleotide m^6^A sites by incorporating site-specific antibody specificity

**DOI:** 10.1101/2024.02.04.578119

**Authors:** Yang Guo, Zehong Wu, Weisheng Cheng, Zhijun Ren, Yixian Cun, Jinkai Wang

## Abstract

Various genome-wide and transcriptome-wide technologies are based on antibodies, however, the specificity of antibodies on different targets has not been characterized or considered in the analyses. The antibody-based MeRIP-seq is the most widely used method to determine the locations of N6-methyladenosine (m^6^A) on RNAs, especially for differential m^6^A analyses. However, the antibody specificities in different RNA regions and their resulting technical biases in differential m^6^A analyses have not been evaluated. Here, we evaluated the m^6^A antibody specificities using 100 pairs of spike-in RNAs with known m^6^A levels at single sites. Based on two replicates with different m^6^A levels on spike-in RNAs, we realized the m^6^A antibody specificities of the m^6^A sites on spike-in RNAs were greatly varied and mainly determined by the surrounding sequences of the m^6^A sites. Moreover, the MeRIP-seq signal fold change is the function of the real difference in m^6^A levels as well as the m^6^A antibody specificity. We then trained a machine learning model to predict the m^6^A antibody specificities of given sequences and predicted the m^6^A specificities of all RNA sequences surrounding the known m^6^A motif DRACH throughout the human transcriptome. Finally, we developed a Hierarchical statistic model for Differential Analysis of m^6^A Sites (HDAMS) by taking advantage of the predicted m^6^A specificities. We found that HDAMS can accurately determine the differentially methylated single-nucleotide m^6^A sites and the output more functionally relevant results. Our study not only provides a powerful tool for differential m^6^A analyses but also provides a methodological framework for other antibody-based studies to incorporate antibody specificities.

## Introduction

N6-methyladenosine (m^6^A) is a prevalent RNA modification on mRNAs as well as non-coding RNAs^1, 2^. It is catalyzed by the methyltransferase complex composed of the core METTL3, METTL14, and WTAP^3, 4^. Differentially methylated patterns may lead to different epigenetic regulations in biological processes. In the past decade, various sequencing techniques have been developed to investigate the m^6^A signals in transcriptome towards different resolutions of epigenetic studies. The first method to identify m^6^A in transcriptome-wide scale is MeRIP-seq^2^ (also known as m^6^A-seq^1^), it uses m^6^A antibody to capture the m^6^A methylated RNA fragments followed by high-throughput sequencing and subsequent identification of *∼*100bp m^6^A peaks. One of the limitations of the original MeRIP-seq is the requirement of as large as *∼*200 micrograms of input RNAs, the later-developed low-input MeRIP-seq reduces the input RNA requirement to *∼*500ng^5^. Of note, the recently developed scm^6^A-seq^6^ and picogram-scale MeRIP–seq (picoMeRIP–seq^7^) can identify m^6^A in a single cell. In addition, by barcoding the RNAs of different samples, the m^6^A-seq2^8^ can identify and quantify m^6^A in a large pool of samples. The rapid development of MeRIP-seq, especially picoMeRIP–seq, indicates that antibody-based identification and comparison of m^6^A is still a promising method in the future.

Another limitation of MeRIP-seq is the lack of single-nucleotide resolution. To obtain single-nucleotide resolution of m^6^A, a series of m^6^A antibody-dependent and independent methods have been developed, such as miCLIP^9^, m^6^A-CLIP/IP^10^, m^6^ACE^11^, m^6^A-REF-seq^12^, MAZTER-seq^13^, DART-seq^14^, m^6^A-lable-seq^15^, m^6^A-SAC-seq^16^, GLORI^17^, eTAM^18^, etc. However, these methods have not been widely used in identifying the differentially methylated m^6^A between different physiological and pathological conditions due to different reasons such as operation difficulties and lack of stoichiometry. Therefore, MeRIP-seq is still the most widely used method to identify the differentially methylated m^6^A between different conditions up to now.

Several computational methods have been developed to identify the differentially methylated m^6^A peaks of MeRIP-seq. For examples, exomePeak^19, 20^ is a multifunctional tool to identify m^6^A methylated peaks and use Poisson generalized linear model to detect differential m^6^A peaks in two conditional groups; MeTDiff^21^ treats m^6^A differential analysis as testing the difference of signal distribution among two sample groups, and used a beta-binomial mixture model to test the significance of differential changes of m^6^A peaks; DRME^22^ utilizes a negative binomial model to imitate the observed IP read counts in MeRIP-seq data and uses the statistic test model in DESeq^23, 24^ to identify differential m^6^A peaks or genes; RADAR^25^ uses a Poisson mixed model to consider the over-dispersion challenge of read counts due to variability of samples in model fitting. The common strategy of these statistic methods is they are univariate test models and ignore the dependence of adjacent peaks in testing. To accommodate the dependence structure of methylation bins or peaks, a multivariate test model-TCQ^26^ is proposed to test the methylated difference of multiple peaks in gene-level simultaneously based on the count vectors of IP and input trials. Although these methods have been proven to yield promising results, a few drawbacks still need to be further considered in m^6^A differential analysis: (1) Non-specific binding of m^6^A antibody has been reported to cause artificial m^6^A peaks in MeRIP-seq^27^, however, the antibody specificity for each m^6^A site is still unknown. In principle, variations of m^6^A antibody specificities across different m^6^A sites may result in biases in the enrichment of m^6^A by antibody, thus biased m^6^A quantifications and determination of the differentially methylated m^6^A. (2) In most studies with the aim to elucidate the detailed roles of m^6^A played in various biological processes and diseases, determination of the exact single-nucleotide m^6^A sites is of urgent need. Therefore, it is of great advantage to identify the differentially methylated single-nucleotide-resolution m^6^A sites using MeRIP-seq.

In this study, we proposed a new framework for differential analysis of m^6^A sites by considering the effects of antibody specificity. In order to evaluate the m^6^A antibody specificity for each m^6^A site, we first synthesized 100 pairs of 338bp DNA sequences through *in vitro* transcription with m^6^ATP to generate 100 pairs of spike-in RNAs with known m^6^A levels at single sites. We then trained an ensemble machine learning model to predict the antibody specificities of regions surrounding the m^6^A motif DRACH throughout the human transcriptome. Finally, by incorporating the m^6^A antibody specificities of different m^6^A sites, we developed a hierarchical statistic model-HDAMS to determine the differentially methylated single-nucleotide m^6^A sites.

## Results

### m^6^A antibody specificities are determined by the sequences around m^6^A sites

To determine the specificities of the m^6^A antibody, we synthesized 100 pairs of spike-in RNAs through *in vitro* transcription using the 100 pairs of 338bp DNA templates with GC content ranging from 0.4 to 0.6. For 100 pairs of DNA templates, one sequence has only one adenosine nucleotide in the middle and is located within the GAC m^6^A motif; the other sequence is the same as this sequence except the adenosine in the middle is replaced by a thymine nucleotide (T), resulting in a sequence without adenosine. Before *in vitro* transcription, each pair of DNAs was mixed at a proportion ranging from 0% to 100% and then an approximately equal amount of all pairs was pooled together. After *in vitro* transcription with m^6^ATP, UTP, CTP, and GTP using the above-pooled DNA templates, we obtained a mixture of spike-in RNAs that can mimic 100 different RNAs with diverse m^6^A stoichiometry of the single m^6^A site in the middle (Fig.1). We synthesized two batches of spike-in RNAs using the same 100 pairs of DNAs but the two sequences of each pair were mixed with totally unrelated proportions in order to generate spike-in RNAs that can mimic two samples with differentially methylated m^6^A sites. We added the two batches of spike-in RNAs in extracted mRNAs of GM12878 cells to perform MeRIP-seq, respectively. We identified 17,850 and 21,354 m^6^A peaks on GM12878 RNAs of the two replicates of MeRIP-seq with different batches of spike-in RNAs, respectively. 13,317 m^6^A peaks (74.6% and 62.4% of the two replicates) of the two replicates were overlapped (Fig.2a). Moreover, the m^6^A ratios of the common peaks were also significantly correlated (*r* = 0.875, *P* = 0; Fig.2b). The common m^6^A peaks of GM12878 RNAs were enriched near stop codons and significantly enriched for the known m^6^A motif, suggesting the MeRIP-seq was successfully performed (Fig.2c, Supplementary Fig.S1a). By separating the reads with A and the counterpart T nucleotides, we found the m^6^A ratios of the spike-in RNAs with A nucleotides in the middle were significantly higher than the spike-in RNAs with the T other than A nucleotides in the middle as well as the negative controls without A nucleotides (Supplementary Fig.S1b), suggesting the m^6^A antibody selects the methylated spike-in RNAs over the unmethylated ones.

**Figure 1.**
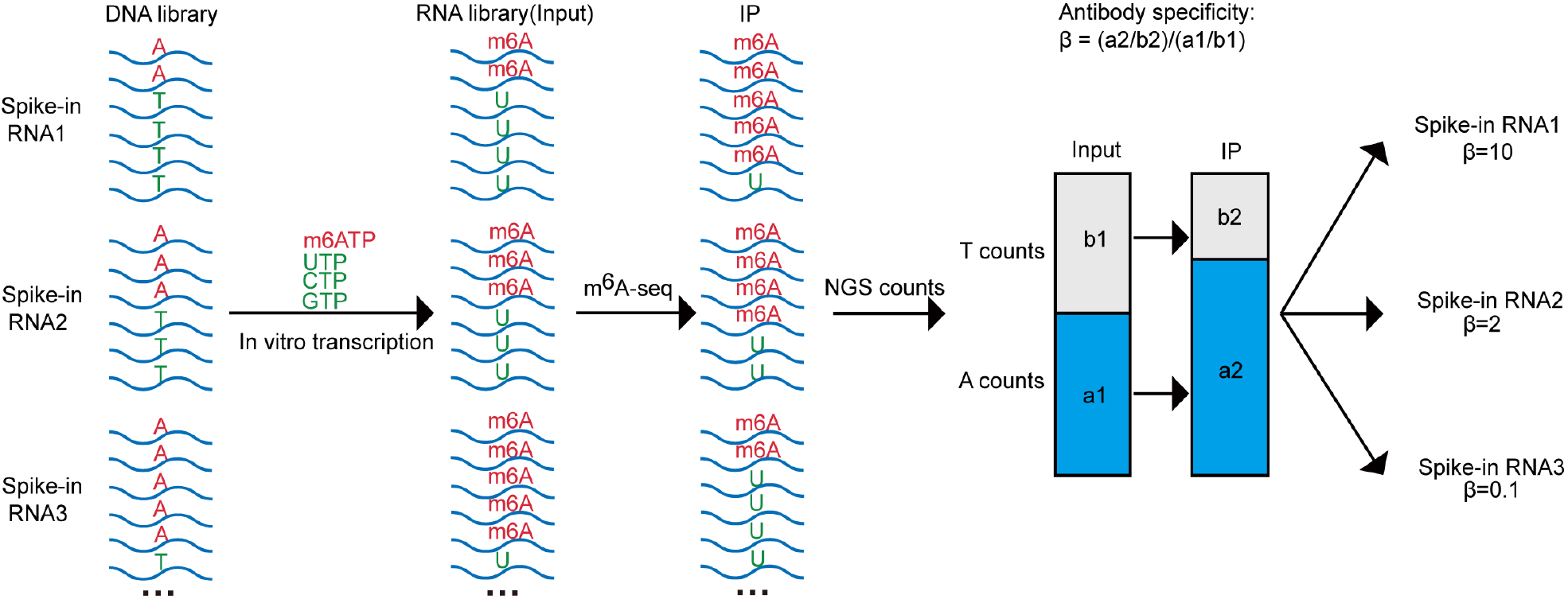
*In vitro* transcription generates spike-in RNA library. Workflow of spike-in RNA synthesis and calculation of antibody specificity. After *in vitro* transcription with m^6^ATP, the A (red) in the spike-ins ultimately yields m^6^A-containing RNA, while the sequences contained a central T (green) paired with it, resulting in RNA without m^6^A. Subsequently, all 100 pairs of spike-ins were mixed into the MeRIP-seq library for experimentation. Three pairs of spike-ins were demonstrated.

Because we converted all the A nucleotides on spike-in DNAs into m^6^A on spike-in RNAs using m^6^ATP other than ATP in the *in vitro* transcription, all adenosines on the spike-in RNAs were m^6^A methylated and all the counterparts in the pairs were unmethylated due to the lack of adenosine. By using the unmethylated counterpart T nucleotide to mimic the unmethylated A nucleotide on spike-in RNAs, the m^6^A level of the m^6^A site can be calculated as the proportion of A reads out of the total of A and the counterpart T reads for each pair of spike-in RNAs. We then calculated the m^6^A antibody specificity (*β*) of the single m^6^A site for each pair of spike-in RNAs as the ratio of the enrichment of A versus the counterpart T (A count / T count) between IP and input (Fig.1, see Methods). As expected, the two batches of spike-in RNAs did have unrelated m^6^A levels based on the calculation (Supplementary Fig.S1c). However, the m^6^A antibody specificities of the m^6^A sites on spike-in RNAs were significantly correlated (*r* = 0.817, *P* = 3.89e-25; Fig.2d), indicating that m^6^A antibody specificity is the function of the RNA sequence other than m^6^A level. On the other hand, we found that the *β* values varied widely, ranging from 0.65 to 22.93 (Fig.2e), with 82% of them were lower than 2, which indicates a 2-fold enrichment of m^6^A-modified RNAs over unmodified RNAs.

**Figure 2.**
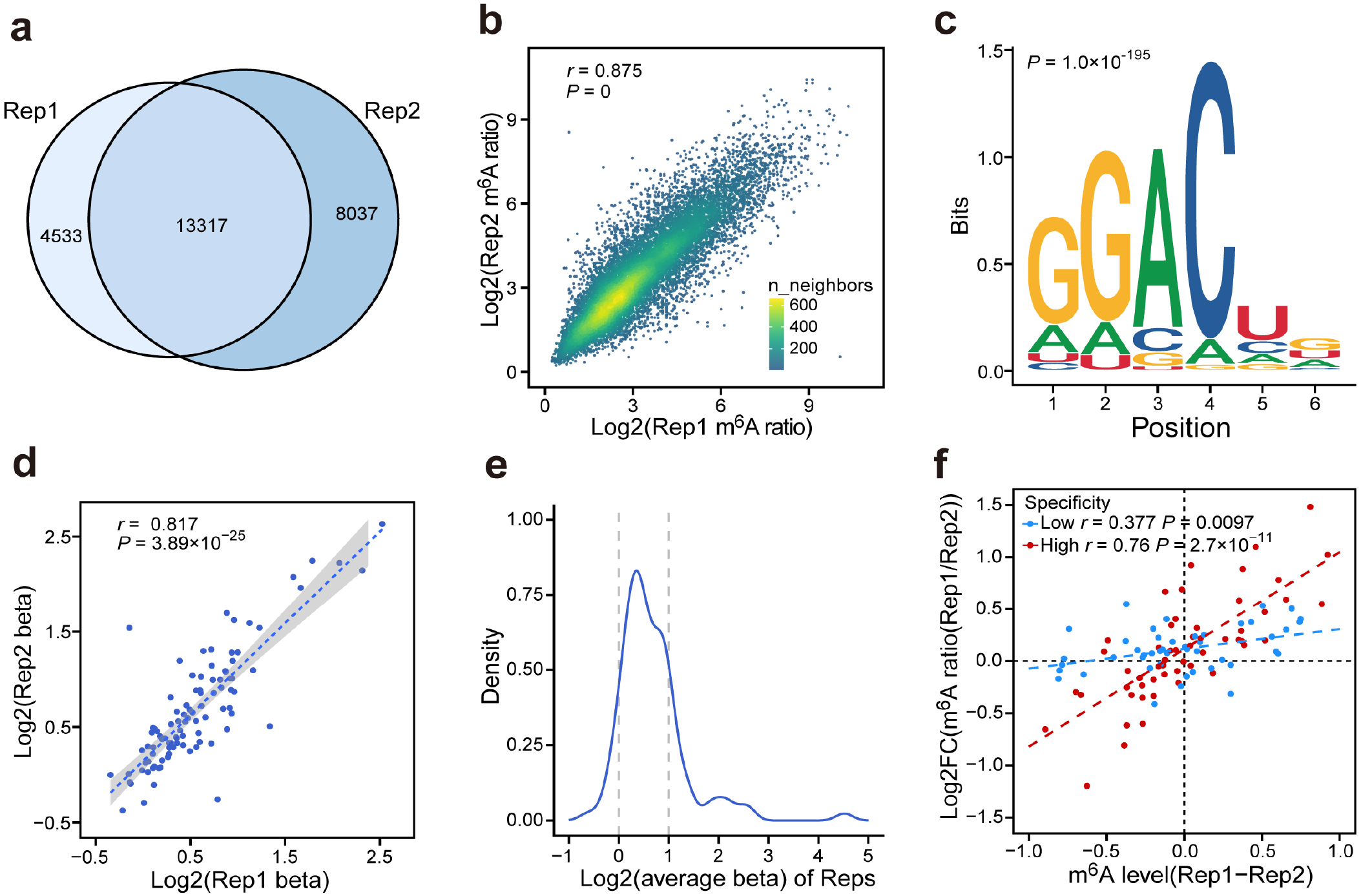
Antibody specificity of spike-in RNAs. **a**. Venn diagram showing the overlap of m^6^A peaks in two biological replicates. **b**. Scatter plot of the m^6^A ratios of common m^6^A peaks between the two replicates (n=13,317). **c**. Motif enrichment of common m^6^A peaks between the two replicates. **d**. The correlation of log2-transformed beta values for spike-ins between two replicates. **e**. The distribution of log2-transformed beta values for the 100 pairs of spike-ins. **f**. Scatter plot of actual m^6^A level differences and detected m^6^A ratio differences for spike-ins between two replicates. Spike-ins were divided into low and high groups based on their beta values, with a threshold set at 1.4 (low n = 46, high n=54). The statistical testing in **b, d** and **f** was performed using the Pearson correlation test.

We then compared the m^6^A ratios, which is the ratio of the read coverages at the m^6^A sites between IP and input, of the m^6^A sites on spike-in RNAs between the two replicates of MeRIP-seq based on the total counts of A nucleotides and the counterpart T nucleotides to mimic the m^6^A quantification in normal MeRIP-seq (see Methods). We found the fold changes of m^6^A ratios between the two replicates were significantly correlated with the expected m^6^A level differences between them, suggesting MeRIP-seq is capable of detecting the m^6^A changes. However, when we divided the spike-in RNAs into two groups with high and low *β* values based on a cutoff of 1.4, we found the slopes of the regression functions of the log-transformed fold changes of m^6^A ratios on the expected m^6^A differences were remarkably different, with 0.934 and 0.190 in the high and low *β* groups, respectively (Fig.2f). This result indicates that MeRIP-seq is much more sensitive in detecting the genuine m^6^A differences on the m^6^A sites with higher m^6^A antibody specificities but lack of power in detecting even large m^6^A difference on the m^6^A sites with low m^6^A antibody specificities.

### An ensemble learning model predicts the m^6^A antibody specificity of each m^6^A motif using the surrounding sequence

Since m^6^A antibody specificity is mainly determined by the sequence around the m^6^A site, we next aim to establish a machine learning model to predict the *β* values using the 75bp sequences flanking each side of the m^6^A sites. As the accurate antibody specificity is hard to predict directly based on limited sequence knowledge, we compromised the problem to predict the probabilities of m^6^A antibody specificity in high and low levels. For this purpose, we designed an ensemble learning model (Fig.3a), which integrates the classifiers of logistic regression (LR) and support vector machine (SVM), to robustly predict the m^6^A antibody specificity of the input 151bp sequences centered around the DRACH motif.

**Figure 3.**
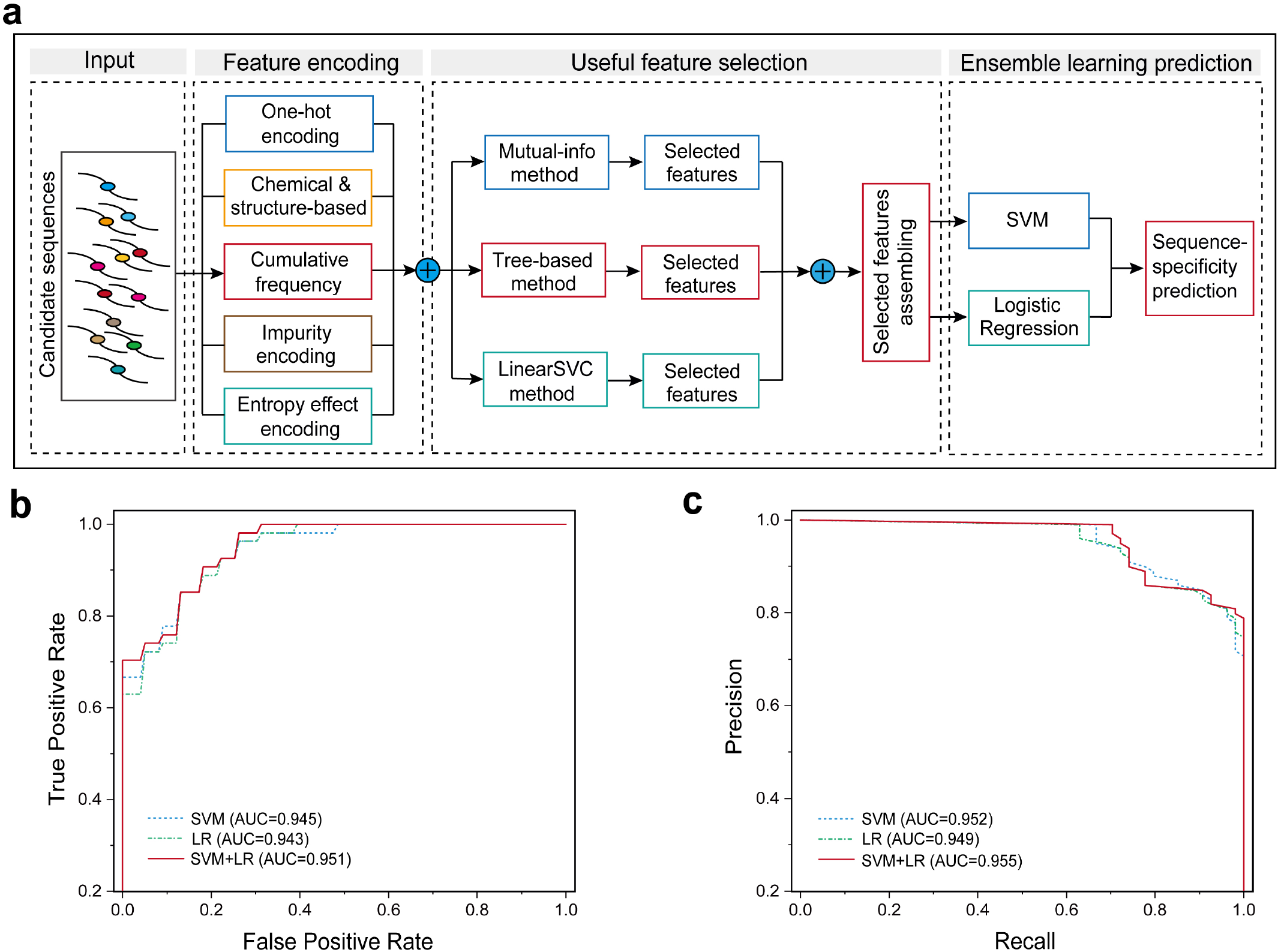
The site-specific m^6^A antibody specificity prediction. **a**. The overall framework of the ensemble learning model for predicting sequence-specificity levels. Five types of sequence encoding methods were used to extract features from multi-views, useful features were selected using multiple feature selection methods, and an ensemble learning model based on SVM and logistic regression classifiers was designed to predict sequence-specificity in low and high beta levels, respectively. **b-c**. The ROC and precision-recall curves of different models to predict sequence-specificity in spike-in data using two-fold cross-validation method.

To predict m^6^A antibody specificity using sequence only, we compiled five types of sequence features. Concretely, we used the one-hot method to encode the position features of nucleotides in sequences and the chemical and structure-based method^28, 29^ to encode the molecule features of nucleotides in sequences, respectively. In order to inspect the local distribution of nucleotides and diversity of local sub-sequences in whole sequences, we also proposed other three types of features to encode the input sequences, including: (1) cumulative frequencies of nucleotides to capture the characteristics of nucleotides’ distribution at different positions in sequences; (2) impurity features to acquire the characteristics of differential distribution of nucleotides in local sub-sequences at different positions versus the whole sequences; (3) entropy effect energy features to measure the information gain characteristics between the sideward local sub-sequences and the central A-included sub-sequence undergoing the penalty of position distance (details in Methods). We used these five types of sequence features extracted from the input sequences as the estimation features in each classifier and trained the ensemble classifying model by using the spike-in sequence data, which was labeled in high and low classes based on the level of known specificity. To assess the prediction performance of the proposed ensemble classification model, we used two-fold cross-validation strategy to evaluate its overall performance on spike-in data. As shown in Fig.3b-c, we found that the ensemble classifying model obtain a superior performance (AUROC=0.951, AUPRC=0.955) and has a better performance than each individual classifier.

### Cellular m^6^A sites with low antibody specificity have less sufficient power in detecting the m^6^A differences

By using the trained ensemble classification model with a prediction probability greater than 0.8, we identified 529,933 and 387,470 DRACH motifs on the exonic regions of the human transcriptome with low and high specificity *β* values based on the 151bp sequences flanking the motifs. Based on the m^6^A levels in HEK293T cells measured by the antibody free method GLORI^17^, we observed no significant difference in the m^6^A levels between high and low specificity of the DRACH sites, consistent with our finding based on spike-in RNAs that antibody specificity is not related to the genuine m^6^A levels (Supplementary Fig.S1d). Compared with the DRACH motifs with high *β* values, the DRACH motifs with low *β* values are overrepresented in protein-coding regions (CDS) (Fig.4a), which is consistent with the previous finding that non-specific m^6^A peaks in MeRIP-seq are enriched in CDS regions^27^. Consistently, the DRACH motifs with low *β* values were significantly enriched in the previously reported non-specific m^6^A peaks as compared with the high-confident m^6^A peaks^27^ (*P* = 4.68e-29, two-tailed Chi-square test; Fig.4b), suggesting that low m^6^A antibody specificity may contribute to the generation of non-specific m^6^A peaks in MeRIP-seq. We then determined 52,778 low specificity and 34,548 high specificity DRACH sites with a prediction probability greater than 0.8 that matched the known single-nucleotide m^6^A sites identified by miCLIP^9^ and GLORI^17^ (see Methods) from diverse human tissues. We found the 151bp regions flanking known m^6^A sites with predicted low *β* values showed significantly higher GC contents (*P* = 0, two-tailed Wilcoxon test; Fig.4c). This observation suggests that the m^6^A antibody may also have a strong binding affinity to the RNAs without m^6^A but with high GC content, which is consistent with the commonly recognized notion that antibodies may prefer to bind the DNA or RNA regions with higher GC content. We then tested whether the *β* value affects the MeRIP-seq based differential m^6^A methylation analyses by comparing the changes in m^6^A during the directed differentiation of human embryonic stem cells (hESCs) toward mesendodermal cells^30^. Consistent with our observations in spike-in RNAs, the annotated m^6^A sites with low *β* values had significantly lower absolute log2-transformed fold changes of m^6^A ratios than those with high *β* values (*P* = 1.09e-12, two-tailed Wilcoxon test; Fig.4d). Consistently, based on our previously calculated coefficient of variations (CVs) of m^6^A ratios across 25 unique cell lines^31^, we found the m^6^A peaks that match the known m^6^A sites with low *β* values had significantly lower CVs (*P* = 1.91e-9, two-tailed Wilcoxon test; Fig.4e), further suggesting the m^6^A sites with low antibody specificity have less sufficient power in detecting the m^6^A differences.

**Figure 4.**
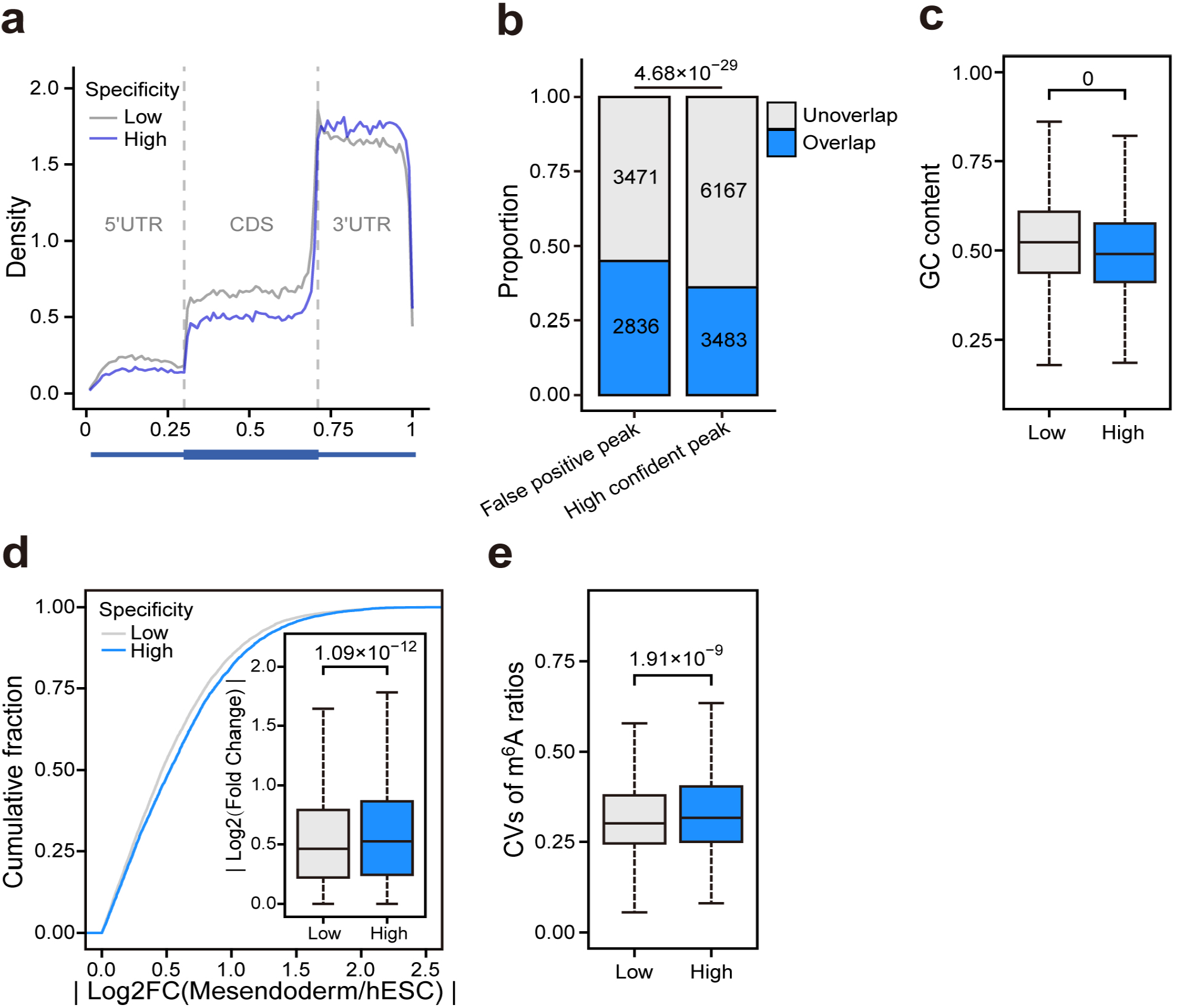
Low specificity m^6^A sites exhibit elevated false positives, and the detection of modification changes is subtle. **a**. Metagene profile showing the normalized distribution of transcriptome-wide DRACH sites with predicted low and high antibody specificity. **b**. The fraction of false-positive and high-confident m^6^A peaks overlap with the predicted low specificity DRACH sites. Due to m^6^A peaks obtained from HEK293T cells, only DRACH sites with RPKM > 10 in HEK293T were used in analysis. Also, peak extension was used to mitigate bias resulting from inherent length differences between the false positive and high confident peaks. Statistical testing was performed using the chi-square test. **c**. Boxplot shows the GC content of 151bp sequences flanking the known m^6^A sites with predicted low and high antibody specificity (low n=52,778, high n=34,548; two-sided Wilcoxon rank-sum test). **d**. The absolute value of log2-transformed fold changes of m^6^A ratios between hESC and Mesendoderm with predicted low and high specificity (low n=11,985, high n=6,465; two-sided Wilcoxon rank-sum test). **e**. The coefficient of variations (CVs) of m^6^A ratios in m^6^A peaks from 25 unique cell lines with predicted low and high specificity (low n=6,349, high n=3,180; two-sided Wilcoxon rank-sum test).

### A statistic model HDAMS accurately identifies the differentially methylated m^6^A sites by incorporating the predicted m^6^A antibody specificities

Because each m^6^A site has its own antibody specificity (*β* value), we then aim to establish a statistic model to integrate the *β* level information to identify the differentially methylated single-nucleotide m^6^A sites from MeRIP-seq based comparisons. For this purpose, we developed a <u>h</u>ierarchical statistic model for <u>d</u>ifferential <u>a</u>nalysis of <u>m</u>^6^A sites (HDAMS) (Fig.5a). In HDAMS, we model the observed counts of each m^6^A motif site from a sampling process, in which the probability of methylation was determined by multiple effect factors. Specifically, we use the negative binomial distribution to model the IP count from the total coverage count of IP and input channels, in which the methylated probability is estimated by a generalized linear model to combine the effect factors of its region-specific antibody specificity, sample group and original m^6^A ratio. Given a candidate m^6^A site *i*, its observing coverage counts in IP and input trials are *Y*_*i j*_ and *X*_*i j*_ on *j*-th sample, respectively; the HDAMS differential model can be described as,

**Figure 5.**
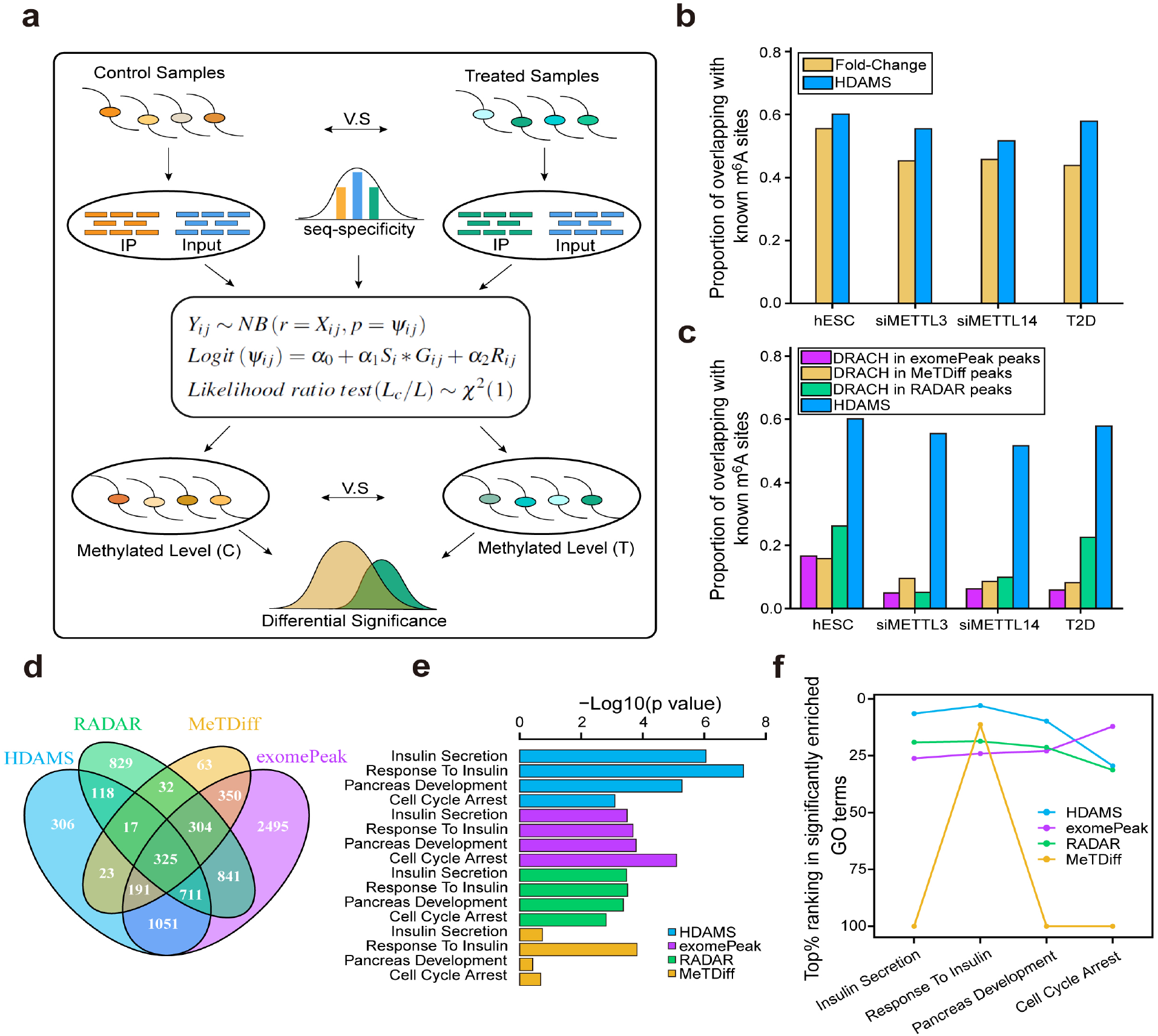
HDAMS outperforms other tools in identifying differential m^6^A. **a**. The diagram of the HDAMS statistic model towards the differential analysis of single-nucleotide m^6^A sites. HDAMS accepts the normalized IP and input counts of samples from different groups and the site-specific m^6^A antibody specificity to test the significance of differentially methylated m^6^A sites in different sample groups. **b**. ===== The performance comparison of HDAMS (FDR<0.05) and Fold-Change (|FC|>1.5) methods in differential analysis of single-nucleotide m^6^A sites in multiple MeRIP-seq datasets. The proportion of overlapping with annotated known m^6^A sites was evaluated. **c**. The performance comparison of HDAMS with three existing methods in the differential analysis of m^6^A sites across multiple MeRIP-seq datasets. The proportion of overlapping with annotated known m^6^A sites was evaluated, and the differential significance of all methods was set as FDR<0.05. As HDAMS focuses on differential single-nucleotide m^6^A sites, the DRACH motif-included sites in differential m^6^A peaks were selected for the other three differential peak-focused methods. **d**. Venn diagram shows the overlaps of genes with differential m^6^A sites/peaks (FDR < 0.05) identified by multiple tools in normal vs. human type 2 diabetes (T2D) islets. **e**. The enrichment of genes containing differential m^6^A sites/peaks (FDR < 0.05, normal vs human type 2 diabetes islets) identified by multiple tools in diabetes-related GO terms. GO analysis was performed by clusterProfiler (v4.0.5). **f**. Top% ranking of diabetes-related GO terms in all significantly enriched GO terms (FDR<0.05). GO terms of genes with differential m^6^A sites/peaks identified by multiple tools were ordered according to FDR. For terms with FDR *≥* 0.05, it was set to the end of significantly enriched GO terms.

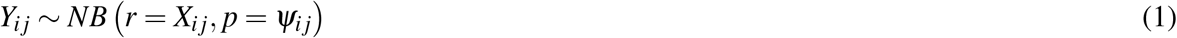

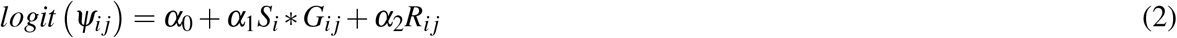

Where *ψ*_*i j*_ is the hidden probability parameter of site *i* is methylated on *j*-th sample; *S*_*i*_ is the probability of site *i* with high antibody specificity level; *G*_*i j*_ *∈ {*1, 2*}* is the sample group label of site *i* on *j*-th sample; *R*_*i j*_ is the original m^6^A methylation ratio of site *i* on *j*-th sample, which is calculated by raw IP and input counts. *α*_0_ is the coefficient of fixed effect; *α*_1_ and *α*_2_ are the coefficients for group and m^6^A ratio effect factors. To test whether there is significant differential m^6^A methylated levels in different sample groups, it can be attributed to the problem of testing whether *α*_1_ is equal to 0 or not. If *α*_1_*≠* 0, then different sample groups will lead to different impacts on the antibody specificity effect trend to the differential response. We use the likelihood ratio test method to estimate the significance of difference between groups. Give the null hypothesis *H*_0_ : *α*_1_ *≠*0; the alternative hypothesis *H*_1_ : *α*_1_*≠* 0; the likelihood ratio between two hypotheses is,

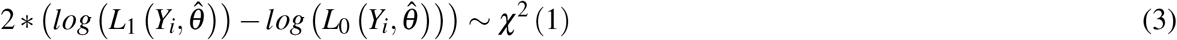

Where *L*_1_ is the maximum likelihood under hypothesis *H*_1_; *L*_0_ is the maximum likelihood under hypothesis *H*_0_; *χ*^2^ (1) is the chi-square distribution with one freedom. By using the above testing model, we can estimate the significance of each site’s m^6^A differential between groups (details in Methods).

### HDAMS accurately identifies differentially methylated single-nucleotide m^6^A sites

We first compared the performance of HDAMS with the traditional fold-change and other three existing popular methods of exomePeak^19, 20^, MeTDiff^21^ and RADAR^25^ using four published differential m^6^A analysis MeRIP-seq datasets, including the m^6^A methyltransferases METTL3 and METTL14 knockdown in human *β* -cell line EndoC-*β* H1 using m^6^A antiboby from NEB^30^, the directed differentiation of hESC toward mesendodermal cells using the m^6^A antibody from Snaptic Systems^32^, and human type 2 diabetes (T2D) islets versus normal using m^6^A antiboby from NEB^30^. First of all, we tested whether the HDAMS improved the performance by comparing it with the method of fold changes in m^6^A ratios at the DRACH motifs. Due to the inherent challenge in directly assessing identified differential m^6^A, we employed an indirect strategy by evaluating the fraction of annotated single-nucleotide m^6^A sites (from miCLIP and GLORI) out of the determined differentially methylated DRACH motifs by different methods. HDAMS identified 1,223, 1,411, 3,022 and 10,026 unique differentially methylated m^6^A sites (FDR<0.05) in these four datasets, respectively. We found that HDAMS consistently outperforms the fold-change method (|FC| >1.5) in all experimental datasets based on m^6^A antibodies from different companies (Fig.5b). Moreover, we further tested whether the HDAMS reported differentially methylated single-nucleotide m^6^A sites are more likely the genuine m^6^A sites than the DRACH motifs in the differentially methylated m^6^A peaks as identified by other commonly used methods, including exomePeak^19, 20^, MeTDiff^21^ and RADAR^25^. We found only a small portion of DRACH motifs in the differentially methylated m^6^A peaks determined by previous methods were known m^6^A sites, in contrast, more than half of the HDAMS determined differentially methylated DRACH motifs are actually known m^6^A sites (Fig.5c), indicating HDAMS has superior power of determining the exact differentially methylated single-nucleotide m^6^A sites. By considering the m^6^A sites identified by miCLIP are also antibody-dependent, to mitigate potential interference from this factor, we conducted the same analysis exclusively utilizing m^6^A sites identified by GLORI, resulting in consistent results with our previous observations (Supplementary Fig.S2a-b).

To test whether HDAMS identified differentially methylated sites are functionally relevant, we took advantage of the MeRIP-seq data of human type 2 diabetes (T2D) islets vs normal^30^, which is a dataset with seven (normal) and eight (T2D) biological replicates and revealed aberrant m^6^A methylation enriched in insulin pathway related genes and is further confirmed by comprehensive functional assays^30^. The previous tools exhibit substantial differences in identifying genes with differentially methylated m^6^A peaks. The proportion of genes identified exclusively by each tool ranges from 4.83% to 39.8%. Metdiff identifies the fewest genes with differentially methylated m^6^A peaks, with the majority (95.2%) being detectable by other tools. In contrast, exomPeak identifies 6,268 genes with differentially methylated m^6^A peaks, many of which (39.8%) are not detectable by any other tools, suggesting a potentially higher rate of false positives (Fig.5d). We then compared the enrichment significance of the genes with differentially methylated m^6^A sites/peaks identified by these tools in diabetes-related gene ontology (GO) terms. We found that HDAMS obtained the most significant p-values across all tools (Fig.5e). Simultaneously, in comparison to other tools, the diabetes-related GO terms identified by HDAMS consistently ranked higher among all significant terms than other tools (Fig.5f). These results strongly indicate that HDAMS provides not only a more precise location of the m^6^A dynamics but also more functionally relevant targets.

## Discussion

MeRIP-seq is currently widely employed for capturing the m^6^A modification states under various conditions. However, existing studies have indicated that differences in the specificity and affinity of m^6^A antibodies limit their ability to detect differential m^6^A^33, 34^. The relatively low specificity of m^6^A antibody resulted in small fold changes of m^6^A ratios in MeRIP-seq partially explains why the dynamics of m^6^A were previously underestimated and whether m^6^A is dynamic is used to be controversial.

While the higher specificity of m^6^A antibody could reflect methylated changes more sensitively in comparisons. This provides a critical clue to accommodate the sequence-specificity of antibody in differential methylation testing. In HDAMS, we used the estimated probability of the site surrounding sequence having high specificity to introduce the rule of antibody specificity effects in different analysis. This is reasonable to emphasize the guide rule of specificity effect warmly. To enhance the weight of the guide rule of specificity effect in differential test, sensitive transformation to the estimated probability of high specificity can be performed before testing to obtain more flexible results at the risk of improving false positive errors. In addition, unlike the previous methods^22, 24^ which used the two-parameter (mean and dispersion) negative binomial model to fit normalized IP count, we used the single parameter form of negative binomial distribution to fit normalized IP count under the reference parameter of normalized input count. It considers both IP and input count information and simplifies the global test model. Actually, it can be further simplified to the discrete binomial model in likelihood ratio test by ignoring the no-effect constant. Although the over-dispersion due to large variability of samples cannot be handled directly, it can be eased by emphasizing the specificity guide rule in testing.

The potential disadvantage of our method is that we do not consider the clustering of m^6^A sites in short regions, because the spike-in RNAs only have single m^6^A site. Previous studies based on single-nucleotide resolution m^6^A identification have consistently reported that a certain fraction of m^6^A sites were located within 100bp of other m^6^A sites^9, 16, 17^. However, these observed clustering of m^6^A sites in gene level does not guarantee they are also clustered on single RNA molecule. Indeed, recent study that identifying m^6^A in single RNA molecule using ONT direct RNA sequencing revealed that m^6^A is randomly distributed on different molecules and very few RNA molecules have multiple m^6^A sites in a cluster^35^. Therefore, the majority of the m^6^A antibody captured RNAs are indeed have single m^6^A site.

In this study, we proposed a new framework to study the sequence specificity of m^6^A antibody and its effects on m^6^A differential analysis. Antibodies are being widely used in diverse large scale technologies such as ChIP-seq and CLIP-seq. The technical effects of m^6^A antibody may also be the case in other antibody-dependent technologies. Because we found antibody specificities are mainly determined by the targeted sequences, the bias of antibody specificity on the detection and quantification of the DNA and RNA targets may cause spurious enrichments of unrelated datasets thus resulting in inauthentic biological explanations. Therefore, although there was no available strategy to systematically evaluate the antibody specificity on different DNA or RNA targets, it is very necessary to be seriously considered in most of the analyses. Fortunately, our study provides an effective strategy to dissect the regional specific antibody specificities, it will benefit not only the studies in m^6^A field, but also all antibody-dependent genome-wide and transcriptome-wide studies.

## Materials and Methods

### Cell culture

GM12878 cells were obtained from GuangZhou Jennio Biotech Co., Ltd., and cultured in RPMI 1640 (Corning) supplemented with 15% FBS (Biological Industries) in T25 tissue culture flask at 37°C with 5% CO_2_. Cells were tested for the absence of mycoplasma contamination using Myco-Blue Mycoplasma Detector (Vazyme).

### Spike-in *in vitro* transcription

The 338bp spike-in sequences were randomly generated with GC content ranging from 0.4 to 0.6, and with GAC m^6^A motif in the middle of the sequence (adenosine positive template, A^+^ templates). The adenosine negative templates (A^*−*^) were same as this sequence of A^+^ templates except the adenosine in the middle is replaced by the one A nucleotide (T), resulting in a sequence without adenosine. The designed DNA templates were synthesis by Genewiz Co., Ltd. with additional *Eco*53KI restriction enzyme sequence (GAGCTC) and T7 promoter (TAATACGACTCACTATAGG) at the 5’ of DNA template, and *Nru*I restriction enzyme sequence (TCGCGA) at the 3’ of DNA template, then All inserted to pUC57 plasmid. Each pair of DNA template plasmids were mixed with same mass and digested using FastDigest *Nru*I (Thermo Scientific), then *in vitro* transcript using the MEGAscript® T7 transcription kit (Invitrogen) with m^6^ATP, UTP, CTP, and GTP, and incubated at 37°C for 2 hours. The products were treated with TURBO DNase (Invitrogen) at 37°C for 15 min to remove the template DNA, and purified using the MEGAclear™ Kit.

### MeRIP-seq

MeRIP-seq was performed based on the protocols previously described by Dan Dominissini et al.^36^ with several modifications. Briefly, total RNA of GM12878 cells was isolated using NucleoSpin® RNA Midi kit (MACHEREY-NAGEL) according to the manufacturer instructions, then poly(A)+ mRNA was selected using Dynabeads mRNA Purification Kit (Invitrogen) and concentrated using RNeasy MinElute Cleanup kit (Qiagen). Purified mRNA mixed with 1% spike-in RNA was fragmented using the 10*×*RNA Fragmentation Buffer (100 mM Tris-HCl, 100 mM ZnCl_2_ in nuclease free H_2_O) and purified with sodium acetate (Sigma-Aldrich), glycogen (Thermo Fisher) and 100% ethanol. 5*μ*g of fragmented mRNA was incubated with 12.5*μ*g anti-m^6^A antibody (Synaptic Systems) in IP buffer (150 mM NaCl, 10 mM Tris–HCl, pH 7.5, 0.1% IGEPAL CA-630, in nuclease free H_2_O) with RNasin for 2 h at 4°C, and further incubated with pre-washed Dynabeads® Protein G for 2h at 4°C. After washing with 1*×*IP buffer (with RNasin) 3 times, the m^6^A enriched fragmented RNAs were eluted from the beads using freshly prepared elution buffer (150 mM NaCl, 10 mM Tris–HCl, pH 7.5, 0.1% IGEPAL CA-630, 6.7 mM of m^6^A, in nuclease free H_2_O) and purified from the beads using RNeasy MinElute Cleanup kit (QIAGEN). Sequencing libraries were generated using the TruSeq Stranded mRNA library prep kit (Illumina). All libraries were sequenced on an Illumina HiSeq2000 platform to produce 40M 150bp paired-end reads.

### Processing of spike-in data

The adapters and low-quality reads from paired-end data were removed using fastp^37^ (v0.22). Subsequently, reads were aligned to the human GRCh37 genome supplemented with designed spike-in sequences using Hisat2^38^ (v2.2.1). Only uniquely mapped reads with alignment scores greater than 30 were used for downstream analysis. For each pair of spike-in sequences, reads covering the central adenine (A) were denoted as Acounts, and those covering the central thymine (T) were denoted as Tcounts. The m^6^A level of each spike-in pair was calculated as Acounts/(Acounts+Tcounts), and the m^6^A ratio for each spike-in pair was determined by the ratio of normalized counts (Acounts+Tcounts) in the IP sample to that in the input sample. Then, we calculated the specificity for each pair of spike-in as Acounts/Tcounts in IP group divided by Acounts/Tcounts in input group according to the spike-in counts data (Fig.1). To sense the relationships between their specificity and sequence characteristics, we divided all spike-in sequences into two levels (high or low) based on the specificity threshold of 1.4. At last, we used these labeled spike-in sequences to train prediction model according to supervised learning, thus to predict the specificity levels of new sequences from transcriptome.

### Processing of MeRIP-seq data

For all MeRIP-seq datasets, adaptors and low quality reads were removed using fastp^37^(v0.22). Reads from IP and input channels were aligned to the reference genome GRCh37 using Hisat2^38^ (v2.2.1). Reads with mapping scores *≥* 30 were used for downstream analysis. In default settings, exomePeak was used for peak calling, and peaks with FDR *≤* 0.05 were considered as confident m^6^A peaks. In GM12878, BEDtools^39^ (v2.29) was used to identify common peaks between two replicates with the key parameters ‘-s’ (require same strand) and ‘-r -f 0.6’ (at least 60% overlap in both replicates). The findMotifsGenome.pl script in Homer^40^ (v4.11) was used to search for the enriched motif in the m^6^A peak region where the random sequences(100bp) on the human genome with matched GC content were used as background sequences for motif discovery. For all candidate m^6^A sites with DRACH motif in transcriptome, the m^6^A ratio of each site was calculated as the IP RPKM divided by the input RPKM of 151bp sequence flanking on the site. We filtered out sites with input RPKM < 10 in each dataset, as well as sites with input RPKM > IP RPKM, to exclude unreliable sites.

### Prediction of sequence specificity of DRACH motif sites using machine learning model

To obtain the antibody specificity of all candidate m^6^A sites, we considered all possible sites in genome that have DRACH (where D: G/A/T, R: G/A, and H: T/A/C) motif in surrounding sequences. For these sites’ sequences, we designed an ensemble learning model to predict each sequence specificity in two levels (low and high). We supposed that the sequence specificity of each candidate site is mainly determined by the sequence characteristics surrounding the central site A, since m^6^A antibody bonds to its target sites is usually affected by the local sequence. In our model, for each candidate site, we obtained the local sequence round its two sides with length of 75bp (in total 151bp) and used the sequence features to predict its antibody specificity level. The overall framework of the proposed ensemble learning model is shown in Fig.3a. In detail, it first extracted the sequence features of each candidate site using multiple types of encoding methods. Then, to reduce the risk of overfitting in model fitting, three feature-selection methods were used to select the useful features from all encoding features. Finally, an ensemble framework that integrated both the support vector machine (SVM) and logistic regression (LR) models was used to predict the site-specific antibody specificity levels using the selected features.

For each site-specific sequence, to inspect various relationships of nucleotides in sequence, we extracted five types of encoding features to sense its local sequence features.

1. **One-hot encoding**: it is one of the most commonly used strategies to encode sentences in natural language processing (NLP) and is also popularly used to encode DNA/RNA sequences. In general, each nucleotide in a sequence is encoded to a 4-dimension binary vector and then all vectors are concatenated to present the computational features of the sequence. In this study, we encoded the sequence in terms of the encoding principle of A to (1, 0, 0, 0), C to (0, 1, 0, 0), G to (0, 0, 1, 0) and T/U to (0, 0, 0, 1), respectively. One-hot encoding features give a simple presentation of the sequential constitution of nucleotides in sequence.
2. **Chemical and structure-based encoding**: as the previous studies^28, 29^, nucleotides in DNA/RNA (A, C, G and T/U) can be classified into different groups in terms of their chemistry and structure features. For example, the adenine and guanine have the chemical constitution with two rings, while the cytosine and thymine have the chemical constitution with only one ring; the adenine and cytosine have similar chemical function properties, while the guanine and thymine have similar chemical function properties; the guanine and cytosine have strong hydrogen bonds in the secondary chemical structure, while the adenine and thymine have relatively weak hydrogen bonds. Similar to the study^28^, to consider these chemical function and structure information of different nucleotides in sequence, we used a 3-dimension vector *c*(*x*_*i*_, *y*_*i*_, *z*_*i*_) to encode each nucleotide in sequence in terms of the following rules,

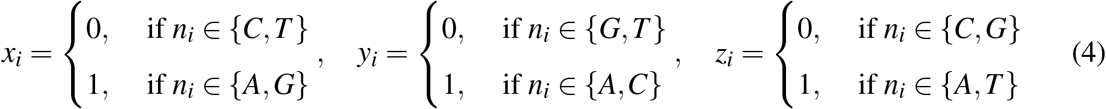

Each nucleotide *n*_*i*_ in a sequence can be mapped to an encoding vector in the form of *c*(*x*_*i*_, *y*_*i*_, *z*_*i*_), likes that A to (1, 1, 1), C to (0, 1, 0), G to (1, 0, 0) and T to (0, 0, 1). The chemical and structure-based encoding features of a sequence can be presented by concatenating all nucleotides’ vectors in the sort of positions in sequence, and they depict the constitution of nucleotides in molecule features of chemical and structure functions.
3. **Cumulative frequencies of nucleotides**: to capture the local distribution of nucleotides in sequence, we proposed a kind of cumulative frequency features of nucleotides at specific positions. Given a sequence *S* with length of *L*-bp, each nucleotide’s cumulative frequency at position *i* is defined as,

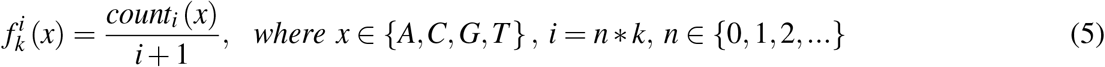

Where 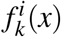is the cumulative frequency of *x* at position *i* in *S* under the skipping parameter of *k*-bp, *count*_*i*_(*x*) is the frequency of *x* in front of the position *i* in *S. k* is the skipping parameter to control the counting size of the feature. For each position *i* in *S* by giving *k*, a cumulative frequency feature of each type of nucleotide can be generated. In this study, we set the pre-defined parameter *k*=3. We finally combined all features of nucleotides in all possible positions to define the features of each sequence *S*. The cumulative frequency features of nucleotides capture the characteristics of nucleotides’ distribution at different positions in sequence.
4. **Impurity encoding**: to consider the characteristics of differential distribution of nucleotides in local sub-sequences at different positions versus the whole sequence, we presented the impurity measure of sub-sequence to define the distance of nucleotide distribution between the local window and background sequences. Given a sequence *S* with length *L* and the window size *W*, a local sub-sequence in the window starting at the position *i* is *w*_*i*_, the impurity of *w*_*i*_ is defined as,

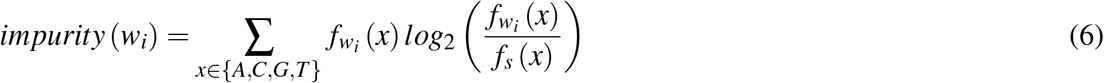

Where, *f*_*wi*_ (*x*) is the frequency of *x* in sub-sequence of *w*_*i*_, *f*_*s*_(*x*) is the frequency of *x* in the sequence of *S*. From the start position of *S*, we defined a series of impurity features by sliding the sub-sequence window with 1bp each time till to reach at the last sub-sequence of *S*. We defined the default value of the pre-parameter *W* =5.
5. **Entropy effect energy encoding**: to inspect the information gain of the sideward sub-sequences to the central A-included sub-sequence, we presented an entropy-based effect energy measurement to encode essential features of the surrounding sub-sequences. For each candidate sequence *S*, whose central site is A with DRACH motif, we defined the central A-included ‘word’ as a sub-sequence with length of *w*-bp at the center of *S*. We suppose the context ‘words’ at both sides of the central ‘word’ in *S* affect the state of central A-included ‘word’. Given a sub-sequence ‘word’ *x* with *w*-bp, the entropy of *x* is defined as,

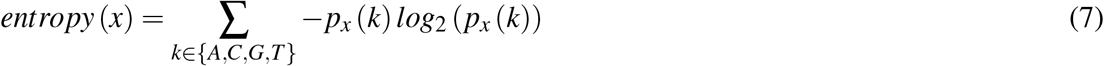

Where *p*_*x*_(*k*) is the probability of nucleotide *k* in word *x*, which is approximated by its frequency in *x*. The entropy of *x* defines the diversity of character information in *x*. To measure the effect of each sideward word *x*_*r*_ to the central A-included word *x*_*c*_ in a sequence *S*, we defined the entropy effect energy measure as,

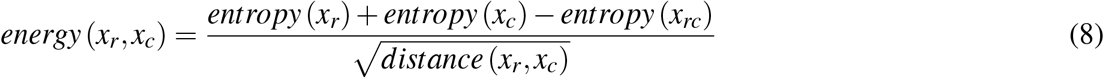

Where *x*_*rc*_ is the joint string of *x*_*r*_ and *x*_*c*_; *distance*(*x*_*r*_, *x*_*c*_) is the gap distance between *x*_*r*_ and *x*_*c*_. The entropy effect energy between two sub-sequence words defines the information variation of the two words in separated and joint forms by considering the distance between them, reflecting the mutual effect of the two words in information gain. Given a candidate sequence *S* and a sliding window *w*, the central sub-sequence word *x*_*c*_ can be easily defined. Then, we defined a series of sideward words (*x*_*r*_) by moving the sliding window 1bp each time in both sides of *x*_*c*_, and calculated the entropy effect energy features between each of them and the central word *x*_*c*_. Finally, we extracted all entropy effect energy features from each qualified position of the raw candidate m^6^A sequence *S*. We used the default value of word length *w*=15 in this study.

- **Feature selection:** for each candidate sequence, we extracted all types of features based on the sequence encoding methods and combined them to define its calculated features. Since the overall encoding features are high-dimension, it is easy to raise over-fitting in model training. To ease this issue, in our model, we used three types of feature selection methods to select more useful features to fit better model. One of the most advantages of using multiple types of methods to select useful features is to consider as more as informative features by enhancing the diversity of features from different views. In detail, we used the k-neighbors mutual information (MI) method^41, 42^, which is a univariate feature selection way based on neighborhoods of the considering feature, to select each type of useful features using classifying strategy. To reduce the variance of estimation, 11-neighbor parameter was set in this study and we finally selected those features with the MI value greater than 0.1. Meanwhile, we also used the linearSVC and tree-based estimators to select useful features in multiple views. The linearSVC (https://scikit-learn.org/) is a selection method which identifies useful features according to using a supervised linear SVM model with penalized L1-norm to obtain sparse solutions. We set the sparse parameter *C* in linearSVC model as 1.0 in default. The tree-based method is another feature selection branch which identifies useful features in classification according to computing impurity-based importance of features based on the forest of decision trees (https://scikit-learn.org). The number of estimator trees was set to 101 in this study. We finally selected all types of useful features by those three methods and combined all three types of selected features to define the input sample features of the downstream prediction models. All three feature selection methods were implemented by using the basic models in scikit-learn package(https://scikit-learn.org/).
- **Ensemble learning model:** based on the selected features of samples, we trained an ensemble learning model to predict the sequence antibody specificities of new sites. To enhance the robustness of the prediction, we designed an ensemble learning model in prediction by integrating the support vector machine (SVM) and logistic regression (LR) models. Specifically, the final estimated sequence specificity level of each sample was set as the average level from the two predictors. To train the ensemble model, we used the pre-designed spike-in sequences which the specificity labels were known by dividing the *β* values into low and high levels. Therefore, the final mission is to train an ensemble classification model to predict the probabilities of each new site for its specificity at the low and high levels. We used the linear kernel and L2-regularization with penalty parameter 1.0 in SVM model, and L2-regularization with penalty parameter 10.0 in LR model. Both of the two models were implemented by using scikit-learn package (https://scikit-learn.org/). To obtain better fitting model in training phase, we used two-fold cross validation method to select the best model by balancing the accuracy of the both classifiers as the final prediction model after multiple trials. Finally, for each new m^6^A site’s sequence, we first obtained its useful features using the methods of feature processing; then, the finally fitted ensemble classification model was used to predict its sequence specificity distribution at low and high levels.

### The hierarchical statistic model for differential analysis of m^6^A sites

Numerous studies indicate that a limited specificity of m^6^A antibodies poses challenges for MeRIP-seq in detecting differential m^6^A modifications under diverse conditions^33, 34^. We also observed that same situation in our spike-in data (Fig.2f). Therefore, it is important to consider the effect of sequence specificity of candidate m^6^A sites in differential analysis. By using the proposed sequence specificity prediction model, it is easy to estimate the sequence specificity of sites by inputting the local circumstance sequences of the candidate m^6^A sites. To consider the sequence specificity of candidate m^6^A sites in differential analysis, in this article, we developed a hierarchical statistic model for differential analysis of m^6^A sites-HDAMS. The overall framework of the proposed HDAMS is shown in Fig.5a.

- **HDAMS statistic model:** consider a candidate m^6^A site *i* in differential analysis, and let *Y*_*i j*_ and *X*_*i j*_ are the numbers of counts on *j*-th sample mapped to IP and input trails, respectively. Similar to the model in^21^, given the total count of IP and input, the IP count *Y*_*i j*_ can be approximately modeled as sampling from the total count by a specific type of discrete distribution with the parameter of real methylation signal. HDAMS is a hierarchical model to incorporate multiple effect factors to depict the sampling process of IP count by using negative binomial distribution under a certain methylation signal parameter. Given the methylation signal *ψ*_*i j*_ of site *i* on *j*-th sample, the statistic model of HDAMS is,

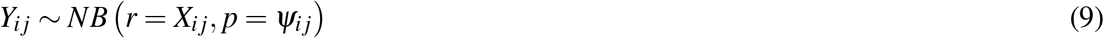

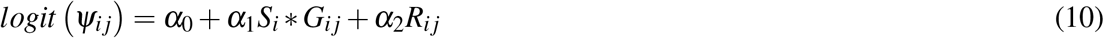

Where *S*_*i*_ is the probability of the predicted sequence specificity of site *i* at high level, which is the differential effect factor of specificity to the methylation signal of the site; *G*_*i j*_ is the group identity of *j*-th sample, and *G*_*i j*_ *∈ {*1, 2*}*; *R*_*i j*_ is the methylation ratio calculated by raw IP and input counts; *α*_*i*_ is the associated coefficient for corresponding factor. Obviously, in HDAMS, we supposed the real methylation signals of a candidate m^6^A site in different samples were directly affected by the sequence specificity, sample group and corresponding methylation ratio factors according to a generalized linear model. The mission of differential analysis of m^6^A site is equivalent to testing against the null hypothesis that *α*_1_=0. As the study in our spike-in data, we found that the trend of differential is more significant at high sequence specificity level (Fig.2f). So, for the sequence specificity effect factor, HDAMS only considers the probability of sequence specificity at high level. Based on the statistic model, given the parameter set *θ*, the likelihood function of the observing IP count data for m^6^A site *i* in *n* samples is,

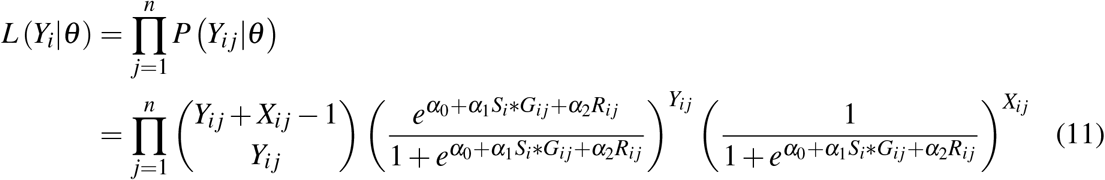

Thus, the logarithm likelihood of observing data vector *Y*_*i*_ can be re-written as,

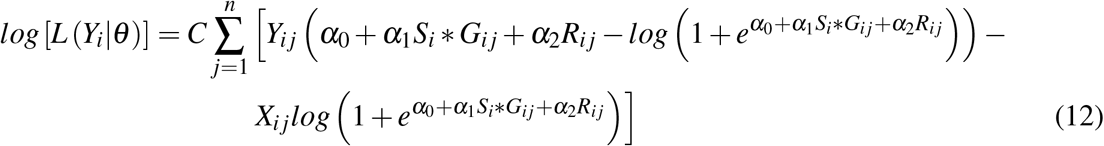

Where *C* is a constant with no effect in calculation. We used the gradient descent algorithm to obtain the maximum likelihood estimators of all parameters. Then, we used the likelihood ratio test method to estimate the significance of methylated differential between groups for each specific site *i*. The null hypothesis and alternative hypothesis of the likelihood ratio test are *H*_0_: *α*_1_ = 0; *H*_1_: *α*_1_*≠*= 0; then, the corresponding likelihood ratio,

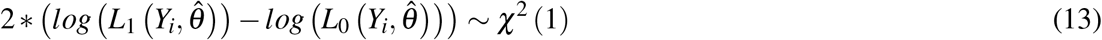

Where 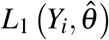is the maximum likelihood of observing vector *Y*_*i*_ under the condition of hypothesis *H*_1_; while, 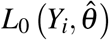is the maximum likelihood of observing vector *Y*_*i*_ under the condition of hypothesis *H*_0_; *χ*^2^ (1) is chi-square distribution with one freedom. Finally, for each candidate m^6^A site, the differential significance of methylation between sample groups can be estimated by the test model. In our pipeline, we used the L-BFGS-B gradient descent algorithm^43^ to estimate all parameters. Notably, if |*α*_1_| *≥ c >* 0, the whole likelihood will be greater than the likelihood on the condition of |*α*_1_| *≤ c*, therefore, to obtain more conservative estimation, we can limit |*α*_1_| *≤ c* in *H*_1_ in practice. We set the default value of *c* as 0.1 in our testing and it can be set as necessary.
- **DRACH sites for differential m^6^A analysis:** we obtained the mapped IP and input counts on the corresponding 151bp sequences of candidate DRACH sites using BEDtools^39^(v2.29) multicov with default parameters. To further obtain more reliable candidate motif sites, we filtered the candidate sites by two criterions: (1) both the average IP count and average input count *≥* 20, and also no input count < 20 in all or most samples; (2) sequence of the site overlaps at least one calling peak *≥* 25bp. The reasons for this selection are to filter the outliers and obtain more reliable sites. We used the exomePeak^19, 20^ method to call m^6^A peaks during the selection of candidate sites for HDAMS analysis.
- **Count data normalization:** to consider the batch effect and sequencing noise of samples in differential analysis, the IP and input counts were normalized separately before using HADMS. We used the same method of median-of-ratio in DESeq2^24^ to perform normalization for count data, as it is robust to both sample and item outliers. Given all *m* samples in analysis, for each site item *i* on *j*-th sample, the raw count number is *K*_*i j*_, then the normalized count 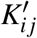 is,

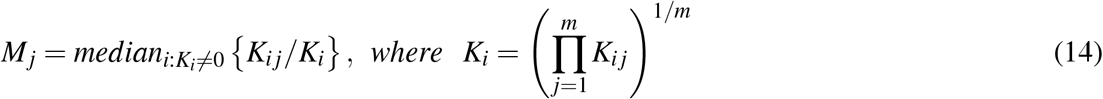

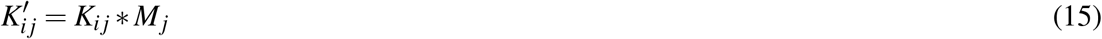

Where *M*_*j*_ is the median of estimated size factor of all items on *j*-th sample. The normalized count 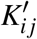 is scaled by the size factor of its corresponding sample.

### MeRIP-seq data in differential analysis

Four MeRIP-seq datasets were used for differential m^6^A analysis, including the disease states (Normal vs. human type 2 diabetes (T2D) islets GSE120024^30^), knockdown conditions (Control vs. METTL3 and METTL14 knockdown in the human *β* -cell line EndoC-*β* H1 GSE132306^30^), and differentiation process (hESC vs. differentiated mesendodermal cells GSE52600^32^). These datasets were utilized to assess the impact of antibody specificity on differential m^6^A analysis and to test the performance of different tools in detecting differential m^6^A sites. Differential m^6^A analysis was performed on each dataset using HDAMS, exomePeak, MetDiff, and RADAR with default parameters. Sites/peaks with FDR < 0.05 were identified as differential methylated m^6^A sites/peaks.

### Collection of single-nucleotide m^6^A sites from miCLIP and GLORI

To evaluate the performance of different tools in m^6^A differential analysis, we collected 122,129 m^6^A sites identified by miCLIP from public NCBI GEO datasets GSE63753^9^, GSE98623^44^, GSE121942^45^, GSE122948^46^, GSE128699^47^, GSE134103^48^. As miCLIP sites lack quantitative m^6^A information, we also collected 170,126 m^6^A sites with m^6^A level identified by antibody-free method GLORI^17^. Finally, we collected a total of 234,051 unique m^6^A sites for filtering unreliable DRACH sites as well as investigating the performance of HDAMS.

### m^6^A peaks across different cell lines

To assess the impact of antibody specificity on m^6^A site across different cell lines, we used m^6^A peaks identified in 25 unique cell lines from our previous research^31^. We calculated the coefficient of variations for m^6^A ratios of m^6^A peaks with low and high antibody specificities across 25 unique cell lines. To assign the m^6^A antibody specificity of each m^6^A peak, the antibody specificity of a known m^6^A site that overlaps with the m^6^A peak was used. In cases where an m^6^A peak contained multiple m^6^A sites, the mean probabilities of low and high antibody specificity were calculated separately. At last, peaks with probability values >0.8 were categorized as either high-specificity or low-specificity m^6^A peaks, while the remains were filtered out.

## Data availability

The raw sequencing data generated in this study has been deposited in the Genome Sequence Archive (GSA) for Human under the accession number HRA006398 (reviewer accessible link: https://ngdc.cncb.ac.cn/gsa-human/s/1Uw4n77A).

## Code availability

The source code of HDAMS is available on GitHub at https://github.com/glabatlas/HDAMS.

## Acknowledgements

This work was supported by the National Natural Science Foundation of China (62102173, Y.G.; 32270630, J.W.; 31970594, J.W.; 32300455, Y.C.), the Guangdong Science and Technology Program (2022A1515110255, Y.C.), the Natural Science Foundation of Gansu Province (21JR7RA509, Y.G.), the Fundamental Research Funds for the Central Universities (23xkjc003, J.W.; 22qntd4810, Y.C.; 23ptpy90, Z.R.).

## Figures & Tables

**Figure S1.**
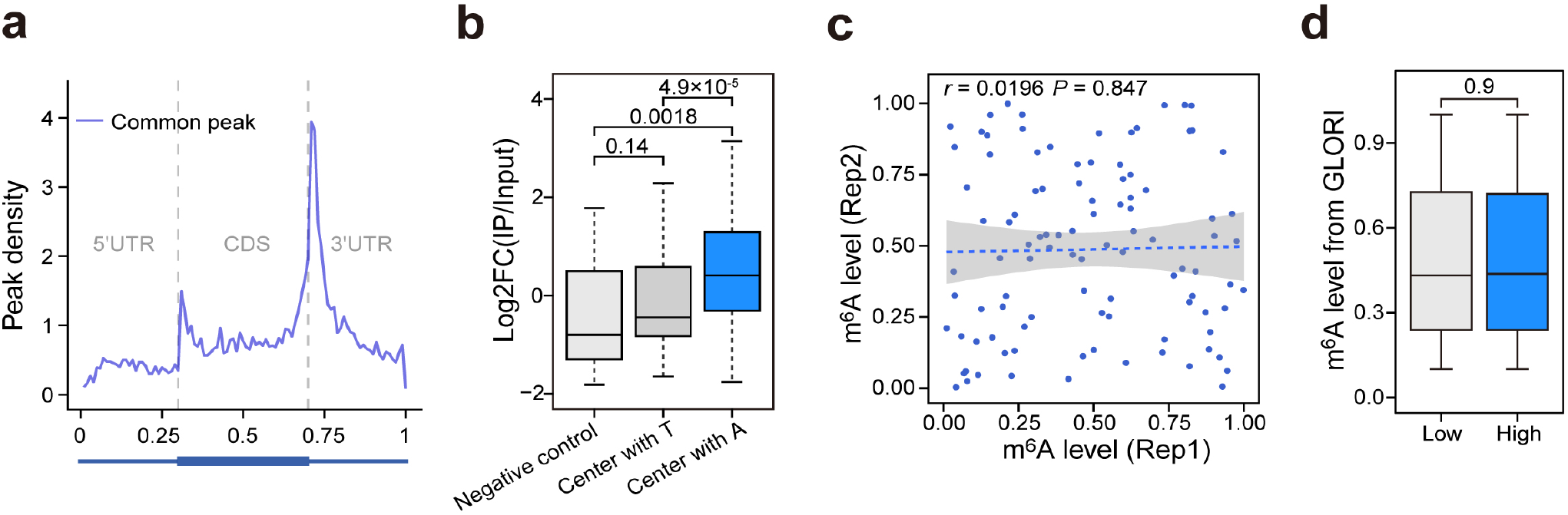
Antibody specificity of spike-in RNAs. **a**. Metagene plot of the normalized distribution of common m^6^A peaks across 5’UTR, CDS, and 3’UTR of mRNAs. **b**. The m^6^A ratios (normalized IP counts/Input counts) of the 100 pairs of spike-ins with A and T centers, as well as 20 negative control sequences without A. **c**. The correlation of the m^6^A level of spike-ins between the two replicates. **d**. The m^6^A level of known m^6^A sites identified by GLORI with predicted low and high specificity (low n=39,266, high n=25,235).

**Figure S2.**
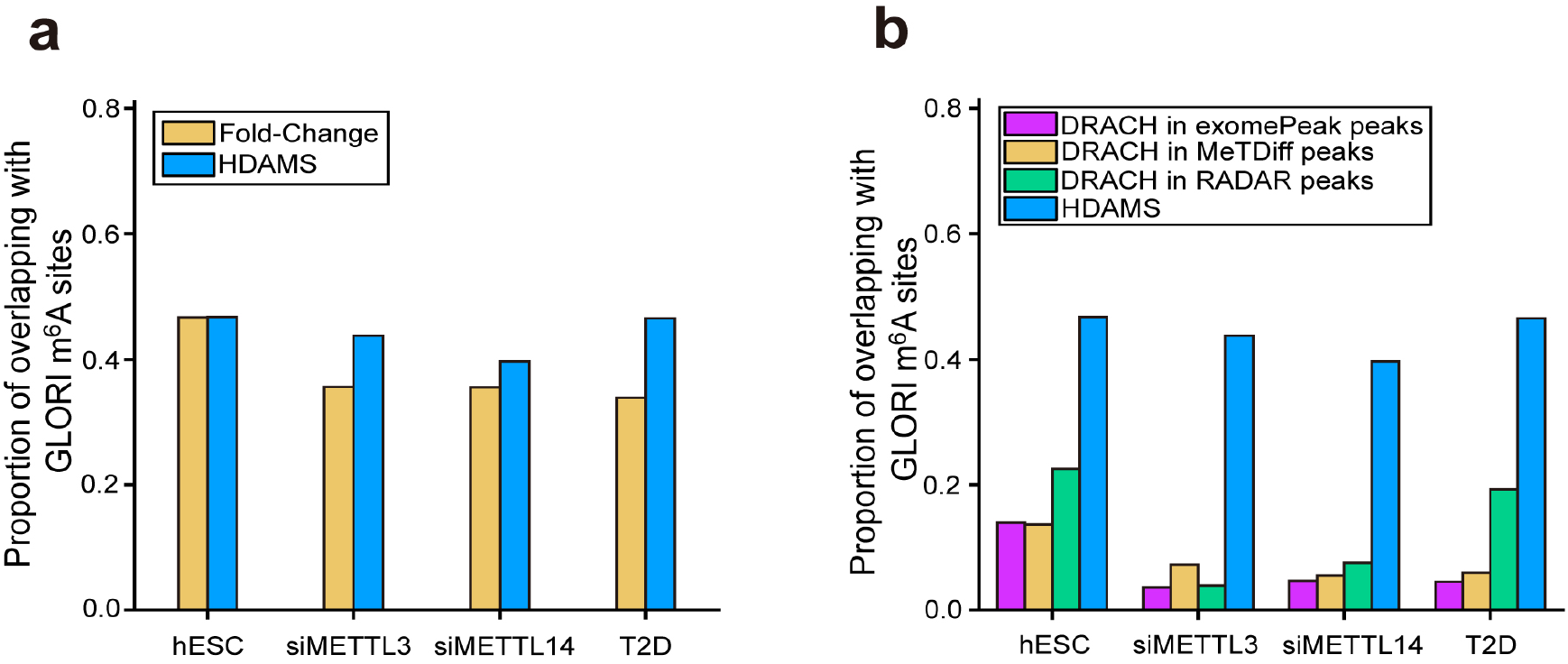
Comparisons of different methods in identifying differential m^6^A sites. **a**. The performance comparison of HDAMS (FDR<0.05) and Fold-Change (|FC|>1.5) methods in differential analysis of single-nucleotide m^6^A sites in multiple MeRIP-seq datasets. The proportion of overlapping with GLORI m^6^A sites was evaluated. **b**. The performance comparison of HDAMS with three existing methods in the differential analysis of m^6^A sites across multiple MeRIP-seq datasets. The proportion of overlapping with GLORI m^6^A sites was evaluated, and the differential significance of all methods was set as FDR<0.05. As HDAMS focuses on differential single-nucleotide m^6^A sites, the DRACH motif-included sites in differential m^6^A peaks were selected for the other three differential peak-focused methods.

